# An improved assembly of the *Albugo candida* Ac2V genome reveals the expansion of the “CCG” class of effectors

**DOI:** 10.1101/2021.03.29.437041

**Authors:** Oliver J. Furzer, Volkan Cevik, Sebastian Fairhead, Kate Bailey, Amey Redkar, Christian Schudoma, Dan MacLean, Eric B. Holub, Jonathan D.G. Jones

## Abstract

*Albugo candida* is an obligate oomycete pathogen that infects many plants in the Brassicaceae family. We re-sequenced the genome of isolate Ac2V using PacBio long reads and constructed an assembly augmented by Illumina reads. The Ac2VPB genome assembly is 10% larger and more contiguous compared to a previous version. Our annotation of the new assembly, aided by RNASeq information, revealed a dramatic 250% expansion (40 to 110) in the CHxC effector class, which we redefined as “CCG” based on motif analysis. This class of effectors consist of arrays of phylogenetically related paralogs residing in gene sparse regions, and shows signatures of positive selection and presence/absence polymorphism. This work provides a resource that allows the dissection of the genomic components underlying *A. candida* adaptation and particularly the role of CCG effectors in virulence and avirulence on different hosts.

## Introduction

Oomycetes of the order Albuginales are obligate biotrophs and cause a white rust disease on plant hosts that resembles disease caused by basidiomycete rust fungi. Leaf pustules disseminate asexual spores that germinate on new leaves to enter the host via stomata and then emerge via formation of new pustules (blisters) beneath the epidermis that rupture at maturity to re-disseminate asexual spores (Holub et al. 1995). A key distinction, however, is that the asexual inoculum of oomycete rusts is a zoosporangium that releases motile spores in water, which swim and alight on stomata to initiate host penetration. Oomycete rusts also have an overwintering sexual phase, producing oospores from fertilization within leaf tissue which are released into soil as the host tissue decomposes, and then re-infect the next cycle of host plants via roots.

Oomycete rusts have evolved as angiosperm pathogens and include the genus *Albugo* which attacks brassica species (*Albugo candida*), spinach (*A. occidentalis*) and sweet potato (*A. ipomoeae*), and *Pustula* and *Wilsoniana* which attack Compositae (e.g, sunflower) or Chenopodiaceae, respectively (Brandenberger et al. 1994; Kamoun et al. 2015; Thines and Spring 2005). Interestingly, *Arabidopsis thaliana* is a natural host of two species, namely *A. laibachii* which commonly occurs on rosette leaves (Thines et al. 2009) and *A. candida* which attacks floral tissues (Fairhead 2016) in wild host populations. Subspecies or phylogenetic races of *A. candida* have co-evolved in different wild and domesticated host species of Brassicaceae (Jouet et al. 2019; Pound and Williams 1963). The species was first described as a pathogen of *Capsella bursa-pastoris* (Shepherds Purse), now referred to as *A. candida* race 4, and is also commonly found on other wild relatives including *A. thaliana, A. lyrata* and *Cardamine spp*. (Holub et al 1995; Jouet et al 2019). Other subspecies that have specialised on economically important vegetable and oilseed brassicas include Race 2 from *B. juncea*, Race 7 from *B. rapa* and Race 9 from *B. oleracea*.

During infection of a compatible host, Albugo grows as a network of aseptate hyphae between mesophyll cells of the host leaf, and then penetrates through cell walls to produce specialised feeding structures called haustoria by invaginations of the host plasma membrane. Haustoria provide an intimate interface for coordinated nutrient acquisition and suppression of host defense responses by delivery of effector proteins. Intracellular nucleotide-binding leucine-rich repeat (NLR) immune receptors in a resistant host enable specific detection of effectors and triggering of innate immunity. Interestingly, a single NLR protein has been reported which confers broad spectrum white rust resistance (*WRR4A*) to *A. candida* races 2, 4, 7 and 9 (Borhan et al. 2008). However, natural pyramiding of multiple NLRs has been proposed to explain the divergence of *A. candida* subspecies as a consequence of host species-level resistance (Cevik et al. 2019).

Genome sequencing revealed that *A. candida* and *A. laibachii* have compact genomes of ∼40 megabases (Mb) that show signatures of obligate biotrophy, for example lacking certain key enzymes and Necrosis and Ethylene inducing peptides (NEPs) commonly present in phytopathogen genomes. A unique class of secreted effector candidates, that possess what was termed the “CHxC” amino-acid motif, can in a motif dependent manner translocate inside plant cells and suppress plant immunity (Kemen et al. 2011; Links et al. 2011). Analysis of variation in five additional *A. candida* genomes representing four races (2, 4, 7, 9) showed that recombination followed by clonal propagation likely underpins the emergence of new strains (McMullan et al. 2015). Pathogen-enrichment sequencing (PenSeq) on the CHxC effector repertoire and a 400 kilobase (kb) region of 91 field samples revealed that host plant species and *A. candida* races were assorted congruently in terms of phylogeny, suggesting that host adaptation and specialization occur in the field. That study also provided evidence that certain *A. candida* races have increased ploidy levels, with the likely outcome that these lineages can only propagate asexually (Jouet et al. 2019).

Genes encoding effector proteins are often amongst the most variable in the genome and can be embedded in repetitive regions that hinder genome assembly (Raffaele et al. 2010). We speculated that available Albugo genome assemblies were incomplete, and likely lacking up to 20% of sequences. To improve our understanding of *A. candida* infection and to gain further insights into *A. candida* effector repertoires, we used long-read sequencing platforms to generate an improved genome assembly of an *A. candida* race 2 strain from Canada (Ac2V). This, combined with new RNA-Seq data, allowed us to generate a more complete annotation, and expand the number of candidate CHxC (now renamed CCG) effectors more than two-fold. Analysis of the CCG repertoire revealed that i) CCGs are polymorphic and show presence/absence variation amongst *A. candida* races, and ii) they have expanded differentially in comparison to the related species *A. laibachii*, which may be driven by pressure to avoid recognition by host immune receptors. Consistent with this, two companion papers (Redkar et al. 2021 and Castel et al. 2021) report the multiple CCG effectors recognized by different alleles of the resistance genes *WRR4A* (Borhan et al. 2008) and *WRR4B* (Cevik et al. 2019). Our analysis of the Ac2V genome provides an essential foundation for further investigation of *Albugo* effectors.

## Results

### Sequencing and assembly of a PacBio based Ac2V reference genome

We extracted high-molecular weight DNA from a Canadian isolate of *A. candida* race 2 (Ac2V) (Rimmer et al 2009) and submitted it for sequencing on the PacBio RSII platform. Before assembly, raw PacBio reads were corrected with previously obtained Illumina reads (McMullan et al. 2015). Following error correction, we obtained reads with a total length of 1,219,162,236 bp (approximately 30.5x coverage) and N50 of 7093 bp. The corrected reads were then used for *de novo* genome assembly. We named the assembly Ac2V “PacBio”, hereafter “Ac2VPB”.

Compared to the Ac2VRR (SOLID-based; (Links et al. 2011)) and AcNc2 (Illumina-based; (McMullan et al. 2015) genomes our 39.9 Mb assembly is longer and more contiguous with an N50 of 466 kilobases (kb) and an average contig size of 196 kb (Table 1, Figure 1A). Ac2VRR has a similar scaffold length distribution, but has a high number of Ns (1.7 Mb), representing gaps in the sequence. Subtracting those and splitting non-contiguous scaffolds reveals the relative contiguity of the Ac2VPB assembly (Figure 1A).

**Table 1.** Genome statistics

|  | Ac2vRR | Ac2vPB |
| --- | --- | --- |
| Scaffolds/Contigs | 252 | 198 |
| N50 size (bp) | 375,021 | 466,138 |
| N50 (scaffold/contig) | 33 | 29 |
| Average contig length (bp) | 137,158 | 196,775 |
| Assembly content (bp) | 34,563,972 | 38,961,604 |
| Gaps (Ns) | 1,728,198 | 0 |
| % Ac2v reads mapping | 78.26 | 95.50 |
| Repeat content (bp) | 5,875,875* | 11,287,808 |

\*excluding Ns

**Figure 1.**
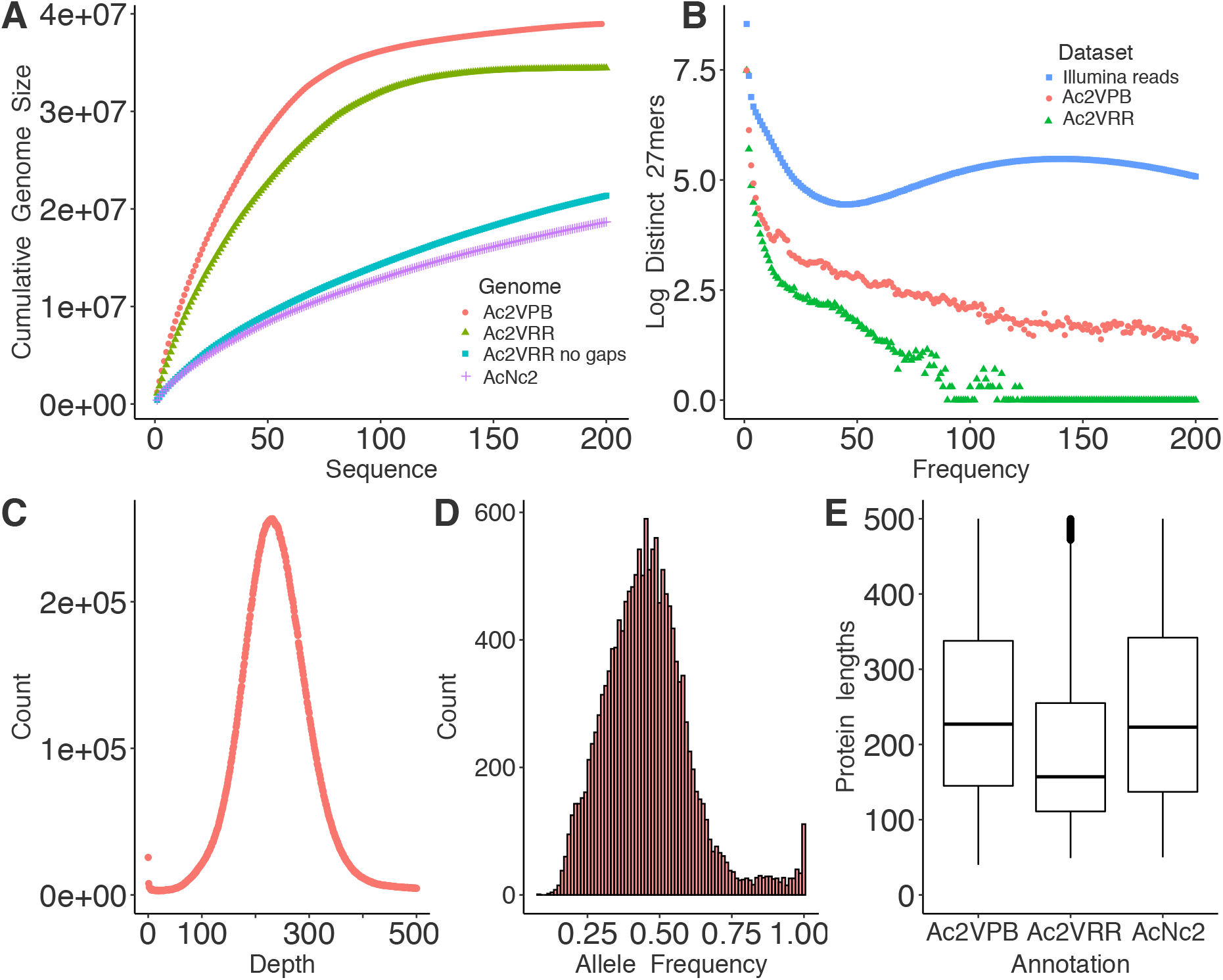
PacBio sequencing produces a more complete *Albugo candida* Ac2V genome and annotation. 1A. Plot comparing the cumulative increase of genome size by size-sorted sequence (contig or scaffold) in the Ac2VPB, Ac2VRR, Ac2VRR ‘no gaps’ and AcNc2 assemblies. The first 200 sequences only are displayed. Ac2VRR ‘no gaps’ and AcNc2 have many further smaller sequences that were thus excluded. 1B. Plot of the log number of unique kmers of length 27 bp (27mers) and their frequency of occurrence in Ac2VPB and Ac2VRR assemblies compared to Ac2V Illumina reads. 1C. Plot of counts of Illumina read depth across the Ac2VPB genome. 1D. Histogram of allele frequency at all sites showing allelic variation when Illumina reads were mapped to the Ac2VPB genome. 1E. Boxplots of the distribution of protein lengths comparing the annotation of Ac2VPB, Ac2VRR and AcNc2.

We aligned Illumina reads from Ac2V to each genome and found that the Ac2VPB genome allowed the mapping of 95.5% of total reads, compared to 78.3% for Ac2VRR (Table 1). The previous Ac2VRR genome was constructed using scaffolding and contained 1.7 million Ns, showing that large parts of the genome were unresolved, likely due to repetitiveness. To compare the repeat content of the genomes, we plotted the frequency of unique 27mers and compared them to the raw Illumina data. This revealed that the Ac2VRR genome is missing many kmers which occurred repeatedly (25-200 occurrences), whereas these are represented in the Av2VPB genome. Comparison to the kmer-content of the Illumina reads (average depth 250) shows that there are some highly repetitive regions (27-mers occurring >300 times) unaccounted for, even in Av2VPB. We estimate the overall repeat content of the Ac2VPB assembly to be 29%, of which half is composed of retroelements (for full analysis of repeats, see Supplemental Data 1).

We further used the mapped Illumina reads to check for either misassembled regions or potential hemizygous regions. Coverage showed a normal distribution centered around 250x and no large region was enriched for either low or high depth (Figure 1C). Likewise, gene depth showed a compact normal distribution, suggesting that most genes are represented at diploid copy number (1 copy, 2 alleles; Supplemental Figure 1A). Of 15,445 quality control-passing SNPs detected, 71% had an allele frequency >0.33 and <0.66, suggesting that Ac2V has a diploid genome (Figure 1D).

### Annotation and search for candidate effector encoding genes

RNA was extracted from Ac2V-infected *Brassica juncea* cultivar “Burgonde” plants at 2, 4, 6 and 8 days post infection (dpi) and used for library preparations and sequencing of 100 bp paired-end reads on Illumina HiSeq2000. The RNASeq reads were then mapped to the Ac2VPB assembly. The overall read alignment rates were 1.8, 17.2, 40.4 and 47 % for 2, 4, 6 and 8 dpi samples, respectively. Trypan-blue staining was used to visualize pathogen growth, which was correlated with the proportion of reads that mapped to Ac2VPB (Supplemental Figure 2).

**Figure 2.**
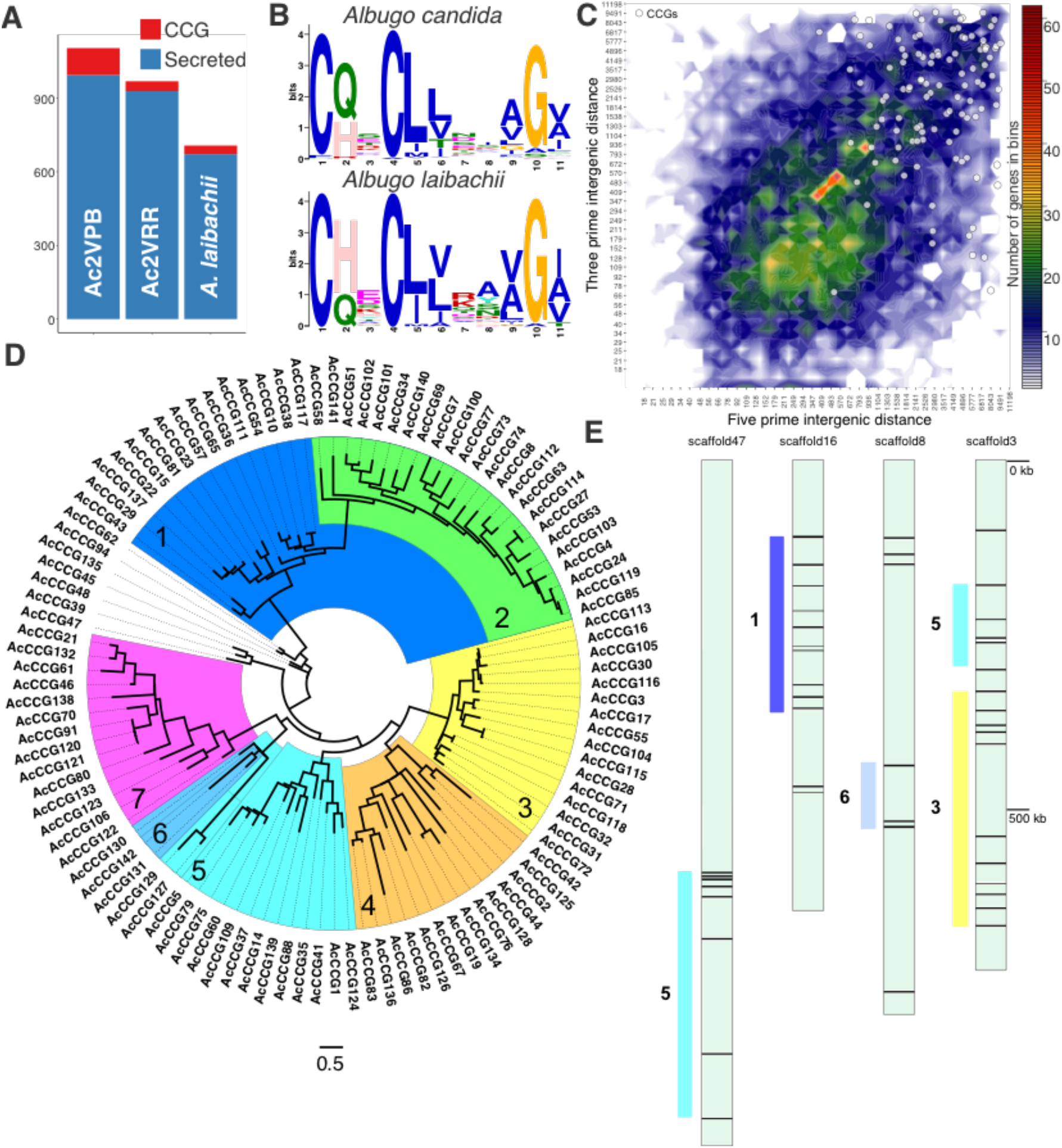
The CCGs are an expanded family of secreted proteins in the *Albugo candida* Ac2V genome. 2A. The CCG motifs in *A. candida* Ac2VPB and *A. laibachii* Nc14. 2B. Barchart of the predicted secretome content of Ac2VPB, Ac2VRR and AlNc14. 2C. Genome density ‘volcano’ plot of the Ac2VPB annotation overlaid with white dots representing the CCG complement. 2D. Maximum-likelihood phylogeny of CCG proteins in Ac2VPB. This is based on an alignment of 60 amino-acids surrounding the conserved CCG motif. Colors and numbers indicate manually annotated phylogenetically related clusters of CCG proteins. Scale bar represents substitutions per site. 2E. Positioning in clusters of CCGs on four selected CCG-rich scaffolds. Colored bars and numbers indicate co-incidence of these clusters with those defined in 2D.

These data were also used as evidence in a holistic gene prediction process (see methods) alongside previous cDNA and protein models from published annotations (Ac2VRR, AcNc2 and AlNc14). The result is a new annotation that contains fewer but longer predicted genes (Figure 1E), and is more complete, as evaluated by BUSCO (Simão et al. 2015) (Ac2VPB: 89% complete fungal BUSCOs vs Ac2VRR: 60% or AcNc2: 82%; Table 2.). The coding space of the new genome is over 1 Mb larger than the Ac2vRR annotation (Table 2.) The AlNc14 genome was reported to lack nitrate and sulphite reductases, and the molybdopterin biosynthesis pathway (Kemen et al. 2011). These are also absent from Ac2VRR and Ac2VPB. Overall, the gene-coding ‘compartment’ of Ac2VPB remains highly compact, with intergenic distances averaging 1.3 kb. The expanded secreted protein complement contained 13 proteins with similarity to CRinkling and Necrosis (CRN) class effectors (Stam et al. 2013) and at least 40 potential cell wall modification enzymes, including 16 candidate secreted glycosyl hydrolases (Supplemental Data 2). A search of all predicted secreted proteins (no transmembrane domain) with the RXLR HMM revealed no RXLR effector candidates (Win et al. 2007).

**Table 2.** Annotation statistics

|  | Ac2vRR | Ac2vPB |
| --- | --- | --- |
| Predicted genes | 15,826 | 13,073 |
| Total protein length | 4,898,153 | 5,869,091 |
| Average protein length | 310 | 449 |
| Predicted secreted proteins without transmembrane domains | 929 | 1104 |
| Complete Fungal BUSCOs number (%) | 173 (59.7) | 250 (89.0) |
| Predicted CCG effectors | 40 | 110 |

We searched the expanded secretome for CHxC effectors. A MEME (Bailey et al. 2009) motif was constructed using a database of CHxC effectors identified from Ac2VRR and AcNc2 and used as input for a MAST search of the Ac2VPB database of proteins with predicted secretion signal (SignalP3.0; (Bendtsen et al. 2004)) and lacking additional predicted transmembrane helices (TMHMM; (Krogh and Rapacki 2016)). From this search we identified 110 CHxC protein candidates, 70 and 75 more than were identified in Ac2VRR and AlNc14 respectively (Figure 2A) and accounting for around 10% of the secretome.

We generated a new motif from the newly expanded CHxC complement. Compared to the consensus motif from AlNc14, we found that the previously reported histidine residue in the CHxC motif was less conserved in Ac2VPB, but a glycine six amino acids after the second cysteine was highly conserved, making the consensus motif CxxCxxxxxG. For simplicity, we redefined the effector class as “CCGs” in *A. candida* (Figure 2B). Within figures in this paper, CHxC names from *A. laibachii* were converted to AlCCG. These names do not indicate orthology between *A. candida* and *A. laibachii* CCGs sharing the same number, CHXC numbers were carried over from Kemen et al (2010).

Amongst the CCGs, only seven showed homology to functionally annotated proteins or domains. Of those 7, 6 were annotated as “AMP-activated protein kinase beta subunit” (Supplemental Data 2). The CCGs were checked for coverage artefacts and show a similar coverage distribution as the overall gene complement (Supplemental Figure 1B).

The CCGs show a significant tendency to have longer than average intergenic distances as revealed by plotting the 5’ and 3’ intergenic distances (Figure 2C) and by comparing combined intergenic distances against other genes (Student’s T-test, p<0.001). We generated an alignment using the CCG and surrounding region of 60 amino acids (the only part where all the proteins in the family could be aligned) and produced a maximum-likelihood phylogeny. This revealed that the CCG effector family in Ac2V falls into 7 major phylogenetic clades (Figure 2D). As a result of the longer contigs of Ac2VPB, it became apparent that these clades largely correspond to physically co-located gene clusters, suggesting that CCGs have undergone parallel segmental duplications forming clusters at different locations across the genome (Figure 2E, Supplemental Data 3).

### Analysis of diversity in candidate effector encoding genes

To assess diversity and selection across the genome we used Illumina sequencing data from six additional *A. candida* isolates of race 4 (AcEx1, AcEm2 and AcNc2), race 7 (Ac7V) and race 9 (AcBoT and AcBoL) (McMullan et al. 2015; Prince et al. 2017). SNP data was used to compare gene-level nucleotide diversity and Tajima’s D ((Tajima 1989); a measure of balancing selection) across the CCGs, predicted secreted proteins and remaining genes. There was no statistical difference between any of these groups (Figure 3A and 3B, ANOVA p-values > 0.05). We noted however the high degree of divergence between these strains which despite being classified as the same species, are differentially adapted to diverse specific host ranges (Jouet et al. 2019); average genome wide identity to Ac2VPB ∼ 98%). This divergence means that in gap-ridden cross-strain alignments of CCG regions, fewer high-quality SNPs could be assigned, resulting in artificially low diversity scores. It was possible to use predicted insertions/deletions (indels) in addition to SNP data to derive an estimate of the proportion of non-synonymous to synonymous changes (pN/pS) and/or pseudogenization in all races in each gene, and in this analysis the CCGs have a significantly higher pN/pS ratio compared to the other two categories (ANOVA, Tukey test p < 0.001, Figure 3C). Further taking into account zero coverage regions, CCGs showed presence/absence polymorphism across the seven races: 27 Ac2V CCGs are absent in one or more of the 6 additional races, and as a class they show a stronger tendency for presence/absence polymorphism compared to other genes or genes encoding non-CCG secreted proteins (Figure 3D), as assessed by the alignment of Illumina reads from the 7 *A. candida* isolates.

**Figure 3.**
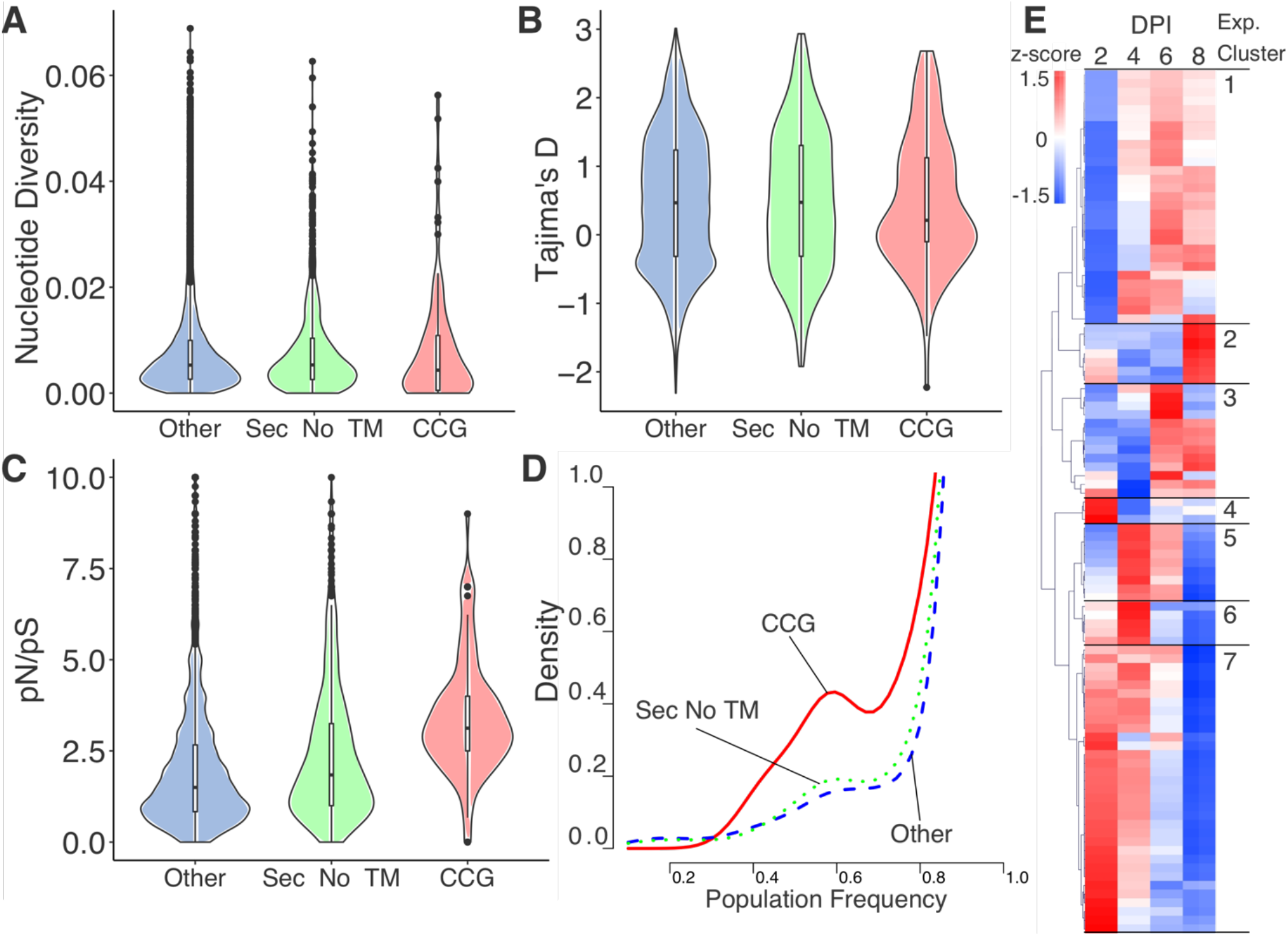
The CCGs in *Albugo candida* are polymorphic and have diverse transcriptional profiles. 3A. Distribution of nucleotide diversity (data from seven *A. candida* races) amongst three classes of genes encoding: CCGs, secreted without additional transmembrane helices and others (the remainder). 3B. Distribution of Tajima’s D (data from 7 isolates) amongst three classes of genes encoding: CCGs, secreted without additional transmembrane helices and others (the remainder). 3C. Distribution of the uncorrected proportion of non-synonymous to synonymous mutations (data from 7 races) amongst three classes of genes: CCGs, secreted without additional transmembrane helices and others (the remainder). 3D. Population frequency of three classes of genes amongst 7 *A. candida* races. Most genes are present across all races (right side of graph). CCGs show an increased proportion of genes present at an intermediate frequency. The y-axis ‘density’ is an arbitrary scale that is necessary to compare density plots across 3 unequally sized groups. 3E. Clustering of CCG encoding genes based on their expression pattern. DPI=days post infection. z-score represents the number of standard deviations away from the normalized mean expression a given measurement is.

The RNASeq reads that were mapped to the assembled Ac2VPB genome were also used to determine the expression levels of CCGs and other secreted protein-encoding genes across all colonization time-points and were grouped into expression clusters (Figure 3E). CCGs showed a full spectrum of expression patterns, from exclusively early expression, to constitutive expression or expression late in the infection. These expression patterns seem to be independent of genomic cluster location, pN/pS ratio or phylogenetic relatedness (Supplemental Data 2).

To compare CCGs in *A. candida* and *A. laibachii*, we constructed a combined phylogeny of these proteins focused around the CCG motif. We found that at the clade level, most CCGs and CHxCs had at least one analog in the sister species, however each had undergone a differential pattern of gene family expansion. Several clusters are greatly expanded in *A. candida* and while others are more expanded in *A. laibachii*. CCGs that have evidence of recognition by WRR4A or WRR4B in Redkar et al (2020) are highlighted and belong to clades that have specifically expanded in Ac2V (Figure 4A). By aligning several of the clades which have expanded in either family, we confirmed that identity is retained in a restricted region around the CCG (Figure 4B, 4C). We observed a frequent feature of two pairs of cysteines located at ∼50 amino acids after the CCG motif. Additionally, we noted complete divergence regarding both identity and length of the C-terminal region occurred in many paralogs (data not shown).

**Figure 4.**
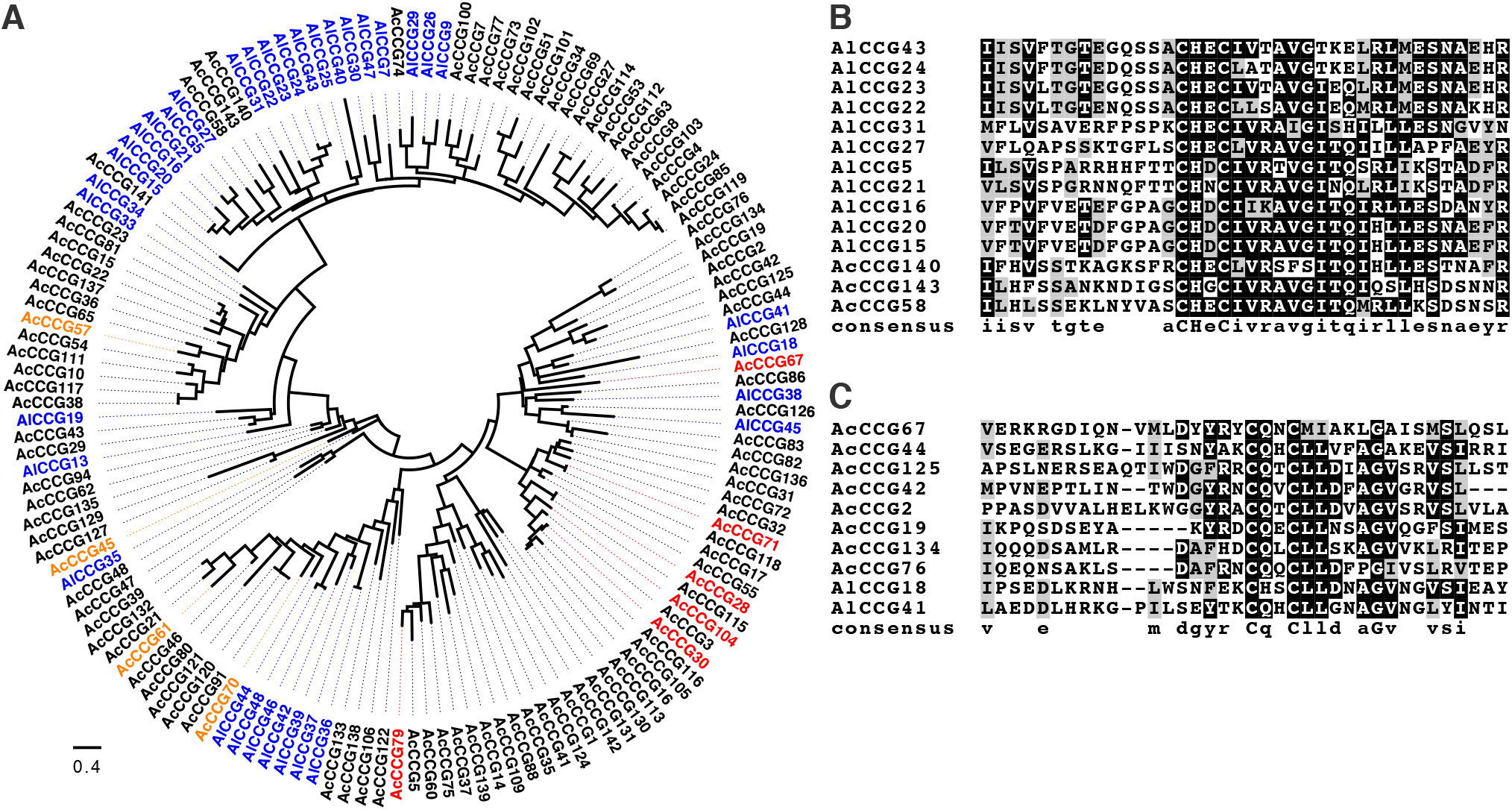
Differential CCG clade expansions in *Albugo candida* and *A. laibachii*. 4A. Maximum-likelihood phylogeny incorporating CCG proteins from Ac2VPB and AlNc14. This is derived from an alignment of the 60 amino-acids surrounding (and including) the conserved CCG motif. Circles indicate the putative Ac2VPB orthologs of CCGs reported recognized by the *Arabidopsis thaliana* Col-0 R proteins WRR4A (red) and WRR4B (orange). Scale bar represents substitutions per site. 4B. Example alignment of an AlNc14 expanded clade. 4C. Example alignment of an Ac2VPB expanded clade.

Based on the expansion of CCGs in the Albuginales, we speculated that ancestral genes might exist in other oomycetes. We investigated CCG presence in four *Phytophthora* species, *Hyaloperonospora arabidopsidis, Pythium ultimum*, and *A. thaliana* as a plant control, using the Motif Alignment &Search Tool (MAST; (Bailey et al. 2009)). We identified 3 potential hits (E-value <0.1) from *P. parasitica* and two from *P. infestans* (Supplemental Data 4). No hits were found in either *P. ultimum* or *A. thaliana*. Only two from *P. parasitica* contained predicted secretion signals directly prior to the CCG motif, a feature of all Albugo CCGs. When inserted into the overall alignment and phylogeny of CCGs from *A. candida* and *A. laibachii*, these candidates form a distant clade with at best 16% amino acid identity in the CCG region to the nearest CCG from Ac2vPB or AlNc14 (Supplemental Figure 3).

## Discussion

We present an improved reference genome for *A. candida*. The genome is still not fully complete, nor assembled into full chromosomes. In the future, methods such as Bionano and Nanopore-based sequencing may enable a fully contiguous *A. candida* genome to be assembled.

The analysis of Ac2V in Jouet et al (2019) was inconclusive about its ploidy level. Non-diploids are generally infertile. In addition to our genome-wide analysis of SNP allele frequency it was reported that Ac2V was capable of mating with an isolate of *A. candida* race 7, with avirulence segregating 3:1 in F_2_ progeny (Adhikari et al. 2003). Together these data support Ac2V as a diploid.

A major outcome of our investigation of the improved genome is the expansion of the predicted repertoire of CCG class effectors by 250%. Other oomycete plant pathogens from the Peronosporalean lineage, which evolved plant pathogenicity independently from the Albuginales, have large families of RxLR and CRN effectors (Baxter et al. 2010; Haas et al. 2009; Torto et al. 2003). The presence of CRN effectors in Ac2V suggests that a common ancestral CRN predates plant pathogenicity, and the expansion to 9 copies may indicate some role in parasitism of plants by *A. candida*.

We performed a phylogenetic analysis of the CCGs and found a link between physical genome location and phylogenetic assortment. The clustering of effectors or virulence factors in microbes is a wide-spread phenomenon, allowing for both the epigenetic co-regulation of these clusters and facilitating mutation and duplication by unequal crossovers (Kämper et al. 2006; Cuomo et al. 2007; Raffaele and Kamoun 2012). This clustering is also seen in fungal pathogens, for example the largest effector gene cluster encodes 24 secreted effectors in the corn smut pathogen *Ustilago maydis* (Brefort et al. 2014). In Ac2V, we found no evidence for transcriptional co-regulation of these clusters, however we propose that the segmental duplication of these CCG clusters occurred through sequential unequal crossovers. Some clusters spanned more than one genome scaffold. Since we have not resolved the genome to chromosome level, we do not know if these are part of the same large cluster on a single chromosome or represent inter-chromosomal transfer of one or more genes followed by further duplications.

In general, gene duplication is considered as a mechanism that facilitates adaptation, as increased gene copy number relaxes purifying selection on each paralog (Ohno 1970); (Soskine and Tawfik 2010). The CCGs have a higher rate of pN/pS and turnover than other genes that encode secreted proteins, suggesting that adaptive evolution of CCG sequence has been key to the co-evolution of *A. candida* races with their respective host species, in line with their predicted role as effectors. Likewise, the tendency of a proportion of CCGs to be absent in various *A. candida* isolates is consistent with these effectors being recognized by host species-specific NLR proteins. Regarding the origin of CCGs, we found a low copy number of distant CCG relatives in other oomycete genomes. More genome sequences of Albuginales and species that lie between them and the Peronosporales are needed to pinpoint the origin of this gene family. Our analysis suggests that following the evolution of a useful ancestor CCG effector, specific expansions occurred as *Albugo* lineages adapted to different hosts. Although we observe the expansion of several distinct CCG clades in *A. laibachii* Nc14, in light of this work it is likely that the CCG repertoire in such Illumina assemblies is under-reported.

Redkar et al (2021) and Castel et al (2021) discovered that certain CCG effectors are recognized by NLRs encoded by *WRR4* and related NLR-encoding genes in *A. thaliana*. Each *A. laibachii* isolate can grow on ∼88% of *A. thaliana* accessions (Kemen et al. 2011), and resistance to *A. laibachii* has not been associated to the *WRR4* locus (Borhan et al. 2004). The pattern of expansion of CHxCs in *A. laibachii* is confined to clades that are distantly related to the recognized CCGs, whereas the expansions in Ac2V are in both the recognized clades, along with several others. We previously proposed that natural pyramiding of NLR-encoding genes such as *WRR4A* and *WRR4B* in *A. thaliana* provides a non-host like protection against races of *A. candida* from brassica hosts (Cevik et al. 2019). We speculate that this has exerted selective pressure on CCG effectors in *A. laibachii* to avoid patterns enriched in CQxC clades, which were free to emerge and expand in for example Brassica-infecting lineages such as *A. candida*. Perhaps a similar NLR-based reciprocal non-host like barrier prevents *A. laibachii* from infecting *A. thaliana* relatives such as *A. lyrata*, and adapted *Albugo* species on those plants have expansions in another variant of the CCG effector class. This is a further reason to obtain high-quality genomes for additional *Albugo* species including ones closely related to *A. candida* (Ploch et al. 2010). Building on this work by obtaining as many diverse samples of CCG family proteins as possible could help unlock the structural basis for CCG recognition, which could lead to the engineering of NLR proteins that confer robust resistance through the recognition of pathogen effector families as opposed to single effector proteins.

Supplemental figures and data files can be downloaded seperately.

Supplemental Figure 1: Gene read depth coverages (aligned Illumina whole genome shotgun reads): a) all genes, b) CCG-encoding genes.

Supplemental Figure 2: Trypan blue staining of Ac2V infection of *B. juncea* at 2,4,6 and 8 dpi, corresponding to RNA-extraction time points.

Supplemental Figure 3: a) Maximum-likelihood phylogeny of CCG proteins from Ac2VPB (orange), AlNc14 (blue) and potential CCGs from *P. parasitica* (strain INRA-310) (black). This is based on an alignment of 30 amino-acids surrounding the conserved CCG motif. Scale bar represents substitutions per site.

Supplemental Data 1: Repeatmasker analysis

Supplemental Data 2: Metadata for all Ac2VPB predicted genes

Supplemental Data 3: Presence/absence and pseudogenization of CCGs across 7 isolates

Supplemental Data 4: CCG MAST hits in proteins with secretion signals

Supplemental Data 6: CCG motif file from MEME

## Author contributions

V.C. and J.D.G.J. conceived the project. V.C. collected the primary data. O.J.F., V.C., S.F., K.B., A.R. and C.S. performed data analysis. E.B.H. provided materials. O.J.F., V.C. and J.D.G.J. wrote the paper. E.B.H., D.M. and J.D.G.J. provided supervision and acquired funding. All authors provided critical feedback during the project.

## Acknowledgements

We acknowledge Ram Krishna Shrestha and Agathe Jouet for technical support and assistance with data retrieval. This research was supported in part by the NBI Computing Infrastructure for Science Group, which provides technical support and maintenance to the Sainsbury Laboratory’s high-performance computing cluster and storage systems.

V.C. was supported by the Biotechnology and Biological Sciences Research Council (BBSRC) Grant BB/L011646/1; O.J.F. was supported by BBSRC Grant BB/M003809/1; A.R. was supported by European Molecular Biology Organization Long-Term Fellowship ALTF-842-2015. K.B. was supported by European Research Council Advanced Investigator Grant 233376 (ALBUGON) (to J.D.G.J.). The Sainsbury Laboratory is supported by the Gatsby Charitable Foundation.

## Materials and methods

### Plant and Pathogen Maintenance

*Brassica juncea* cultivar Burgonde plants were grown on Scotts Levington F2 compost (Scotts, Ipswich, UK) in a controlled environment room (CER) at 22 °C with a 10 h day and a 14 h night photoperiod, and was used for used as host for *A. candida* infections. To propagate *A. candida* race Ac2V, zoosporangia were suspended in cold water and incubated on ice for 30 minutes. The spore suspension was then sprayed on four-week-old *B. juncea* plants. The infected plants were kept under 10 hrs light and 14 hrs dark cycles with a 21°C day and 14°C night temperature.

### DNA Extraction and Genome Sequencing

Zoosporangia were collected from heavily infected *B. juncea*. The spores were then ground to fine powder in pre-chilled pestle and mortar with liquid nitrogen. DNA extraction was then carried out using ChargeSwitch™ gDNA Plant Kit (Invitrogen, CA, USA) following manufacturer’s instructions. *A. candida* Ac2V genomic DNA was sequenced by PacBio RSII platform (4 SMRTcells) (Pacific Biosciences, CA, USA) at Earlham Institute, Norwich, UK.

### RNA Extraction and Sequencing

Four week old *B. juncea* plants were sprayed with the pathogen and between 10-15 infected leaves were collected at 2, 4, 6 and 8 days post infection (dpi) and immediately flash frozen in liquid nitrogen and stored at -80 °C. RNA extraction was carried out using Direct-zol RNA Miniprep Kit (Zymo Research, Cambridge, UK). RNA integrity was then assessed using Agilent 2100 Bioanalyzer with the RNA 6000 Nano Assay Kit (Agilent Technologies, CA, USA). Library preparation was carried out using Illumina TruSeq RNA sample preparation kit (Illumina, CA, USA). Library preparations were then sequenced on an Illumina HiSeq2000 platform and 100 bp paired-end reads were generated at Earlham Institute, Norwich, UK.

#### PacBio Read Correction and Genome Assembly

Prior to the genome assembly, Proovread (Hackl et al. 2014) error correction tool was used to correct PacBio SMRT reads (FastQ) using short Illumina reads from Ac2V (ENA: SRR1811471) (McMullan et al 2015). The genome assembly was then conducted using the error-corrected SMRT reads with Canu V1.3 (Koren et al. 2017).

### Repeat analysis

Repeats were modeled and detected using the RepeatMasker/RepeatModeler pipeline (Smit, Hubley, and Green 2015) including RECON (version 1.08) and RepeatScout (version1.06).

### Annotation

The new Ac2V RNASeq reads from all time points were aligned to the Ac2VPB assembly using HISAT2 (Kim, Langmead, and Salzberg 2015). Successfully mapped reds were extracted and assembled using 3 systems: velvet/oases ((Schulz et al. 2012); versions 1.2.10 and 0.2, kmer sizes 33, 41, 49, 55), soapdenovo ((Luo et al. 2012); version 1.03, trans, kmer sizes 31, 41, 51, 61, 71) and trinity ((Grabherr et al. 2011); version 2.0.6, genome-guided with aligned reads). The resulting redundant pool of transcripts was reduced to the best isoforms using evidentialgene ((Gilbert 2013); tr2aacds v.2013.03.11). These transcripts were aligned to the genome with exonerate ((Slater and Birney 2005); v2.2.0) which provided evidence to predict genes using genemarkES suite v4.21 (Lomsadze, Burns and Borodovsky, 2014). The assembled transcripts were also used as evidence to predict genes using webaugustus v3.2.1 (Hoff and Stanke 2013). Nc2 transcripts and proteins, cloned CCG proteins and transcripts from the Ac2VRR genome were all used to separately train augustus and produce gene models. Independently, all mapped RNA-Seq reads were used to predict genes using braker v (Hoff et al. 2019). All of the resulting gene models, and the exonerate transcript mapping data were together fed into evidencemodeler ((Haas et al. 2008); version 1.1.1) over 3 iterations, which generated a maximal non-redundant genome annotation. A small number of known CCG-encoding genes were manually edited, those genes are annotated with an “m” at the end of their gene or transcript name.

### Gene functional annotation

Eggnog emapper (Huerta-Cepas et al. 2019); version emapper-1.0.3-35) with automatic taxonomic scope was used to assign functional annotations to the Ac2VPB predicted proteins.

### kmer analysis

27mers were counted using kat (Mapleson et al. 2017) and plotted in RStudio using ggplot2 v3.2.1.

### Read alignment and variant calls

Illumina read alignment for whole genome data was performed using BWA v0.7.17 (Li and Durbin 2009) and Samtools v1.10 (Li et al. 2009) was used to process alignments and generate variant calls. VCFtools v0.1.15 (Danecek et al. 2011) was used for VCF formatting and quality filtering (variants with minimum quality of 20 were considered).

### Presence/absence variation analysis

The BEDtools v2.29 (Quinlan 2014) command ‘bedtools coverage’ was used to compute both per base coverage in each alignment, and per gene coverage. Since any % cutoff would be arbitrary, the % coverage of each gene was used to contribute to an overall population frequency score scaled from 0-1 used in Fig 3D. Fig 3D was generated using the sm package in RStudio v1.2.5001 (Allaire 2012).

### Diversity and selection analysis

Nucleotide diversity and Tajima’s D statistics were calculated using popgenome v2.7.1 (Pfeifer et al. 2014) with filtered combined variant call files and the Ac2VPB annotation as inputs.

### Simple pN/pS analysis

Filtered combined variant call files were used as input for SNPEff v4.3t (Cingolani et al. 2012), which divided variants by their computed effect synonymous or non-synonymous to produce a ratio for each gene.

### *ANOVA and Tukey* test

Statistical comparison of population statistics of groups of genes was performed by one-way ANOVA and post hoc Tukey test using the online resource astatsa.com.

### Secreted protein prediction

Ac2VPB proteins were submitted to Signalp3 ((Bendtsen et al. 2004)) and considered secreted. TMHMM v2.0 ((Krogh and Rapacki 2016)) was used to search these proteins post their predicted signal peptide cleavage site for additional transmembrane helices.

### CCG searches

CHXC/CCGs identified by previous genome projects (Links et al. 2011; McMullan et al. 2015; Jouet et al. 2019) were used as a base to generate a CCG motif with MEME ((Bailey et al. 2009); v5.1.1). This motif was used to search the Ac2VPB secreted no transmembrane domain proteins with MAST (Bailey et al. 2009). The proteins positive (E-value <0.1, correct positioning) for the CCG motif were then fed back into MEME to produce a refined CCG motif corresponding to the Ac2V CCG signature. Certain proteins outside the *Albugo* genus, with relaxed E-value threshold < 0.2 are included in the list of hits.

### *Gene Expression* Analysis

For RNASeq data analysis, reads (2 x 100 bp) obtained from each time point were first trimmed using Trimmomatic version 0.36 (Bolger, Lohse, and Usadel 2014) and the quality of the trimmed reads was assessed with FastQC v0.11.4 (Andrews 2014). Reads were then mapped to the assembled Ac2VPB genome using HISAT2 v2.2 (Kim, Langmead, and Salzberg 2015). The counts of reads that mapped to each predicted gene were obtained using the featureCounts utility of the subread package (Liao, Smyth, and Shi 2014). Read count data were then normalized as counts per million (CPM) with EdgeR package (Robinson, McCarthy, and Smyth 2010). Cluster and heatmap were then made using z-scores obtained from normalized (log2(CPM+1)) data.

### Genome architecture analysis

Genome architecture analysis was performed in Rstudio following the protocol of (Saunders et al. 2014).

### CCG phylogenies

CCG proteins were aligned using Muscle and alignments were trimmed and realigned to good quality. These alignments were used to generate maximum likelihood phylogenies using the WAG method, with freq, 3 distinct gamma categories, and 100 bootstraps. These analyses were performed in the MEGA X suite (Kumar et al. 2018). Phylogenies were edited using FigTree (Rambaut 2009).

### Karyoplot

The Karyoplot diagram was generated using karyoplotR (Gel and Serra 2017) using the procedure described in (Van de Weyer et al. 2019).

### General tools for figure production

Figures 1-3 and S1 were generated using the patchwork tool for Rstudio (cran.r-project.org/web/packages/patchwork/) and all figures were edited in Inkscape 0.92 (inkscape.org). ggplot2 v3.2.1 was used to generate scatter plots, box and whisker diagrams and bar charts.

### Data submissions and sources

Data from previous studies, Illumina reads: AcBoT (ENA: SRR1811472), AcEm2 (SRR1806791), AcBoL (SRR1811474), AcNc2 (SRR1811450), Ac2V (SRR1811471). ESTs from Ac2vRR (downloaded from NCBI genbank: HO914811-HO965058, HO965059-HO999999, and HS000001-HS003763), assemblies and annotations of AcNc2, AlNc14, and Ac2vRR.

Data submitted: Ac2vPB PacBio reads (PRJEB39673), Ac2v RNASeq reads, Ac2VPB assembly (GCA_905220665.1), AcEx1 Illumina reads (ERR4395362) Ac7V Illumina reads (ERR5168241), Ac2VPB annotation. The Ac2VPB annotation can be downloaded as GFF or FASTA files at https://github.com/oliverjf/ac2v_genomics.

