## Supplemental_Figures for "An improved assembly of the *Albugo candida* Ac2V genome reveals the expansion of the “CCG” class of effectors"

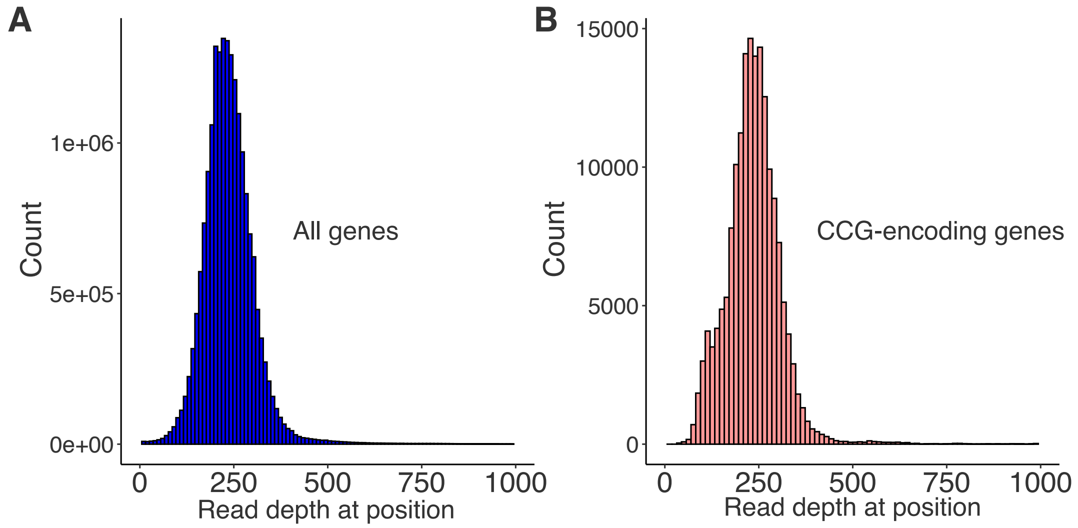


Supplemental Figure 1: Gene read depth coverages (aligned Illumina whole genome shotgun reads): a) all genes, b) CCG-encoding genes.

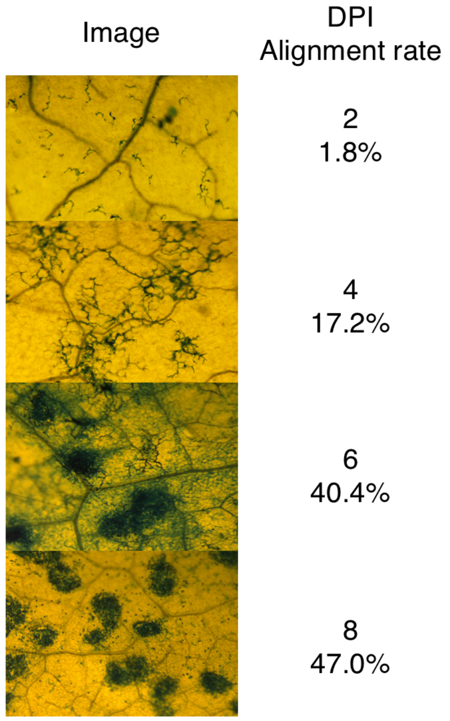


Supplemental Figure 2: Trypan blue staining of Ac2V infection of *B. juncea* at 2,4,6 and 8 dpi, corresponding to RNA-extraction time points.

**A**

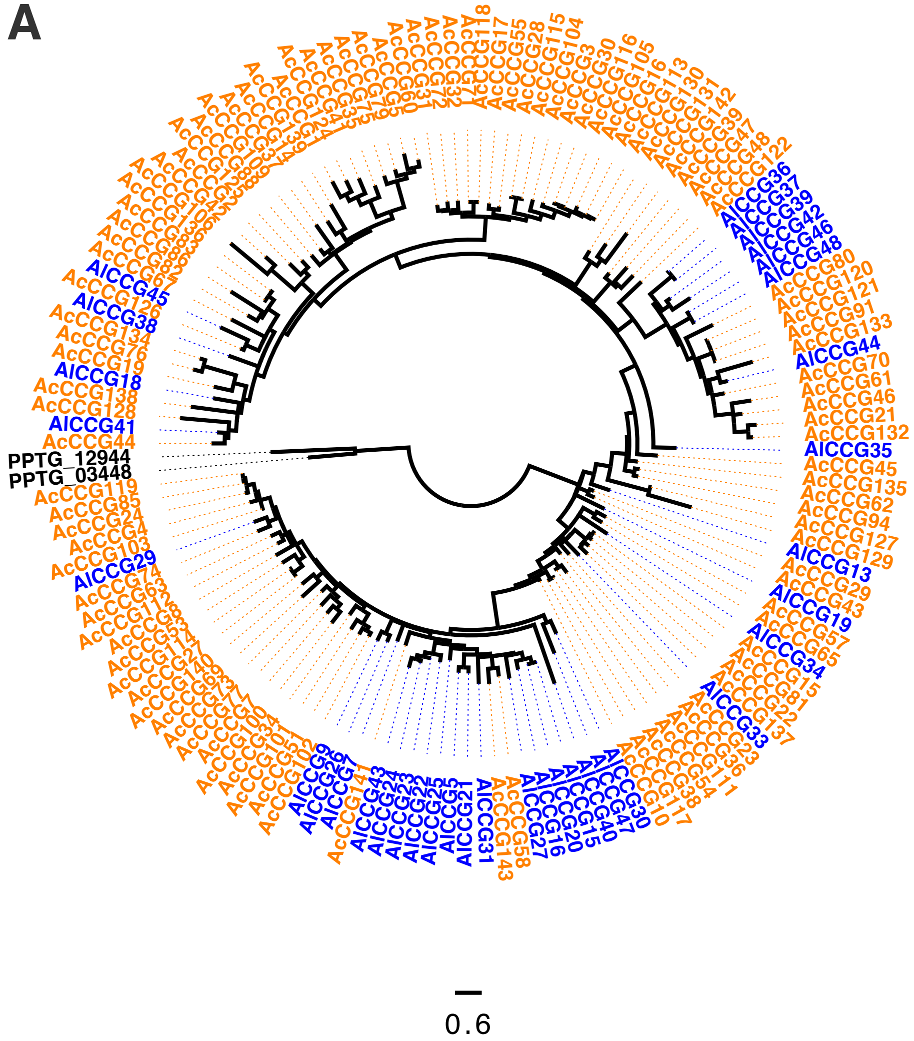


Supplemental Figure 3: a) Maximum-likelihood phylogeny of CCG proteins from Ac2VPB (orange), AlNc14 (blue) and potential CCGs from *P. parasitica* (strain INRA-310) (black). This is based on an alignment of 30 amino-acids surrounding the conserved CCG motif. Scale bar represents substitutions per site.
